# Sevoflurane Increases Locomotion Activity in Mice

**DOI:** 10.1101/447029

**Authors:** Hoai T. Ton, Lei Yang, Zhongcong Xie

## Abstract

Clinical observation shows emergence agitation and hyperactivity during the induction and/or recovery of anesthesia. However, an animal model to illustrate this clinical phenomenon has not been established. We therefore set out to investigate whether sevoflurane, a commonly used anesthetic, could alter locomotion in the mice during the induction and recovery of anesthesia. The activity of mouse was recorded 5 minutes before, during (for 30 minutes) and 40 minutes after the administration of anesthetic sevoflurane [1-, 1.5- and 2-fold minimum alveolar concentration] at 37° C. The total walking distance and velocity of movement were measured and quantified as the indexes of locomotion. We found that the anesthetic sevoflurane increased the locomotion of the mice during the induction of the anesthesia. During the recovery phase after anesthesia, the mice exhibited increased locomotion for a short period of time (about 5 minutes) and then displayed a sharp decrease in mobility for up to 60 minutes following the end of anesthesia administration. The anesthetic sevoflurane did not significantly alter the food intake and body weight of the mice. Furthermore, we found that Alzheimer’ s disease transgenic mice exhibited a greater sevoflurane-induced hyperactivity than the wild-type mice did. Our results showed that inhalation of the anesthetic sevoflurane induced an acute hyperactivity in mice, particularly among Alzheimer’ s disease transgenic mice. These findings from the pilot studies established an animal model to promote further studies into postoperative emergence agitation, hyperactivity and the underlying mechanisms of these conditions.

## Introduction

Sevoflurane is one of the most commonly used general anesthetics. Despite its merits, several clinical studies have reported the emergence agitation and hyperactivity after the administration of sevoflurane, particularly for pediatric patients who exhibit the incidence of emergence agitation up to 80%^1–3^. Moreover, the transient state of marked irritation and disassociation was observed during the transition from unconsciousness to complete wakefulness or following discontinuation of anesthesia in humans^3,4^. Specifically, clinical investigations have shown that sevoflurane can induce hyperactivity during mask induction, even causing body movements, epileptiform electroencephalographic activity, seizure-like movements and actual seizures^1,2,4–7^. Such events pose a risk of injury and potential postoperative complications. However, the causes, underlying mechanisms and targeted intervention(s) of such clinical observations remain unknown.

Recent *in vivo* studies have showed that, particularly during the early stages of AD, neuronal circuits are hyperactive instead of hypoactive^8–11^, which could lead to key changes in AD^12^. Interestingly, sevoflurane has been shown to promote AD-associated apoptosis, Tau protein phosphorylation^10–11^ and β-amyloid protein accumulation (Aβ)^9^, the major pathological hallmarks of AD neuropathogenesis, in cultured cells and in animals. Previous studies have assessed the effects of sevoflurane on neurotoxicity in wild-type (WT) and AD transgenic (Tg) mice^13,14^. However, the acute behavioral change in locomotion following the administration of the anesthetic sevoflurane has not been fully elucidated. Therefore, the objective of the present study was to establish a system to assess the acute effects of sevoflurane on locomotion in mice. The hypothesis in the current studies was that the anesthetic sevoflurane increased locomotion in mice. We further compared the effects of the sevoflurane-induced changes between the WT and AD Tg mice.

## Methods

### Animals

Animal use was conducted in accordance with the National Institute of Health guidelines and regulations. The Massachusetts General Hospital Standing Committee on the Use of Animals in Research and Teaching (Boston, Massachusetts) has approved the animal protocol. Animals were kept in a temperature-controlled (22–23°C) room under a 12-h light/dark period (light on at 7:00 AM); standard mouse food and water were available ad libitum. Housing was provided with appropriate tactile, olfactory, visual, and auditory stimuli. Every effort was made to minimize the number of mice that were used in experiments. The study was performed on WT C57BL/6J and AD Tg 9 month-old female mice (Jackson Lab, Bar Harbor, ME). The AD Tg mice were purchased from Jackson Lab (B6SJL Tg APPSwFlLon,PSEN1*M146L*L286V)6799Vas/Mmjax; Stock Number: 006554), and maintained in our own laboratory. Standard genotyping techniques were used to confirm the condition of AD Tg mice. All efforts were made to minimize suffering of the animals.

### Anesthesia in mice

Prior to carrying out the experiments, we familiarized the mice to the transparent gas-tight plastic chamber (20 × 20 × 15 cm) by placing each of them, without anesthesia, into the chamber for 10 minutes once per day for 3 consecutive days. The experiments were conducted between 9:00 a.m. and 12:00 p.m. Then, the mice were placed into the chamber and exposed to sevoflurane for 30 minutes (for locomotion activity assessment), for 4-hours (for food intake and gain weight assessment) or to the control condition. We used air containing 40% O_2_/60% N_2_ as a carrier, the total gas flow was 2 1/min. During the exposure to anesthesia, the mice were kept warm on a plate heated to 37°C with a white colored background. Control and experimental animals were exposed to the same environmental conditions except that the control animals were exposed only to air. The Dash 4000 monitor (GE Medical Systems information technologies INC., Wisconsion, USA) was used to determine the concentrations of sevoflurane.

### Locomotion measurement

While their horizontal ambulatory activity was measured with the aid of a video-tracking system, the animals were allowed to move freely in the transparent gas-tight plastic chamber. This allowed us to measure all the required parameters offline in simple blind conditions using ImageJ Plugin. The mouse activity was recorded before (5 minutes in air as baseline tracks), during (30 minutes) and after the sevoflurane anesthesia (40-minutes recovery). Here, we used 1, 1.5- and 2-fold minimum alveolar concentrations (MAC), which correspond to 2, 3, and 4% of sevoflurane respectively at 37° C^1,3,15^. The total walking distance (in cm) and velocity (in cm/min) of movement were measured and quantified as indexes of hyperactivity.

Following the experimental session, the mice were carefully removed from the chamber, and returned to their home cage. The test equipment was cleaned with 50% ethanol solution and dried between subjects in order to avoid olfactory cuing. A sample size of 4-9 mice was used for each group and all the mice were used only once.

### Food intake and body weight

The food intake and body weight were measured after daily anesthesia exposure between the hours of 9:00 a.m. and 12:00 p.m. Food and body weight were weighed from days 1 to 4 post-anesthesia. Food intake was calculated as the amount of food removed from the feeding bowl.

### Statistical analysis

Data are presented as mean values with standard error of the mean. The number of samples varied from 14 to 16 in each group for the measurement of food intake and weight changes. The number of mice used for measuring locomotion ranged from 3 to 7 in each group. Data analysis was conducted using the Origin 8.0 software (OriginLab Corporation, Northampton, MA). A student’ s t-test was used to determine the difference between the mice in anesthesia and control condition in terms of the walking distance and velocity. A one-way ANOVA was used to analyze the difference in food intake and weight change between mice in the control group and mice in the anesthesia group. Statistical significance was determined as p<0.05.

## Results

### Sevoflurane induced hyperactivity in mice

Sevoflurane has been reported to induce hyperactivity during the time of anesthesia induction and recovery from the anesthesia in patients^2^. To assess whether sevoflurane would induce hyperactivity in animals, we tracked the horizontal ambulatory activity before and during the induction of the sevoflurane anesthesia as well as after the sevoflurane anesthesia in each of the mice. The total walking distance (in cm) and velocity (in cm/min) of movement were measured offline in a simple blind condition using ImageJ Plugin. As shown in figure 1, all the mice tested had similar baseline levels of velocity ranging from 100 to 130 cm/minute. However, the sevoflurane anesthesia markedly increased the locomotion during the time of the anesthesia induction compared with the control condition as evidenced by the 76% increase in the velocity (194.42 ± 47.23 versus 110.33 ± 21.64 cm/min) and 45% increase in the walking distance (447.95 ± 58.26 versus 309.25 ± 49.25 cm). During the recovery time of the anesthesia, however, there were short periods (within 5 minutes) of increased locomotion, which was followed by decreases in locomotion during the next 20 minutes in terms of velocity (45.86 ± 14.38 versus 99.29 ± 15.52 cm/min) and walking distance (165.91 ± 49.26 versus 270.91 ± 39.26 cm). The activity of mice recovered to the baseline level 60 minutes after the end of the sevoflurane anesthesia (data not shown).

**Figure 1.**
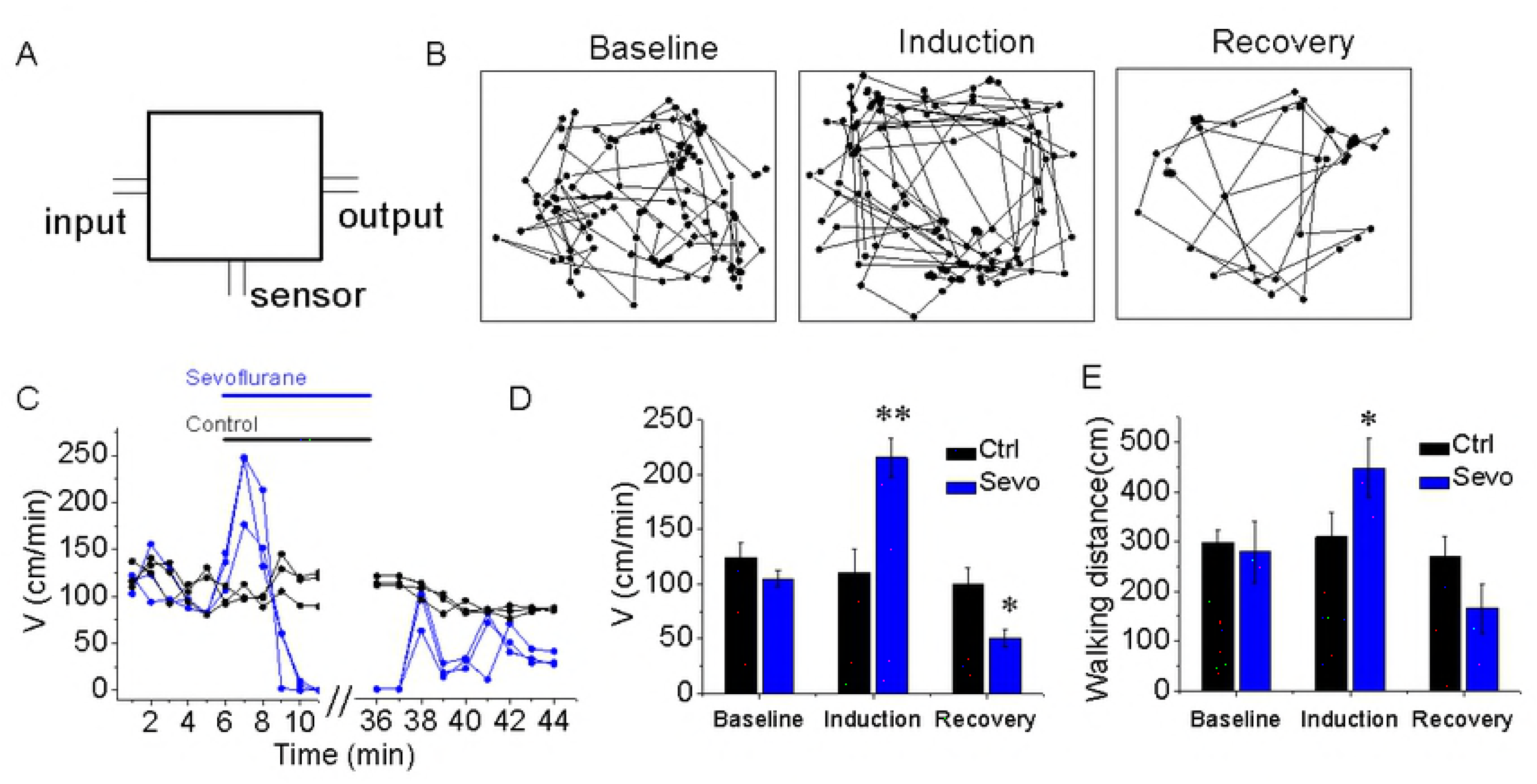
Sevoflurane increases locomotion in mice. (A) Diagram of the chamber used to treat the mice with sevoflurane (2%) in N_2_/O_2_ and access locomotion. Control mice received only N_2_/O_2_. (B) Representative traces demonstrating the position of the mouse in the activity chamber before, during and after administration of 2% sevoflurane anesthesia (C) the time-course of velocity during a single exposure of sevoflurane or control condition to the mice. (D) Summary of mean velocity and walking distance in 2 minutes before, during and after the sevoflurane-treated mice (green bar) and control condition mice (black bar). Data shown as mean and SEM, n = 4-6 in each group.

Next, we tested the effect of sevoflurane on the locomotion at different concentrations ranging from 2, 3 to 4 % (1-, 1.5- and 2-fold MAC). Figure 2 shows that sevoflurane increased locomotion at all of the concentration levels tested. However, we did not observe a statistically significant difference among the different concentrations of sevoflurane on the hyperactivity in the induction phase or hypoactivity in recovery phase of the anesthesia (Figure 2). We noted that the higher concentrations of sevoflurane the mice received, the faster they fell asleep, shortening the duration of their hyperactivity and potentially preventing the possible concentration-dependent effect of sevoflurane on hyperactivity observed during the induction of the anesthesia.

**Figure 2.**
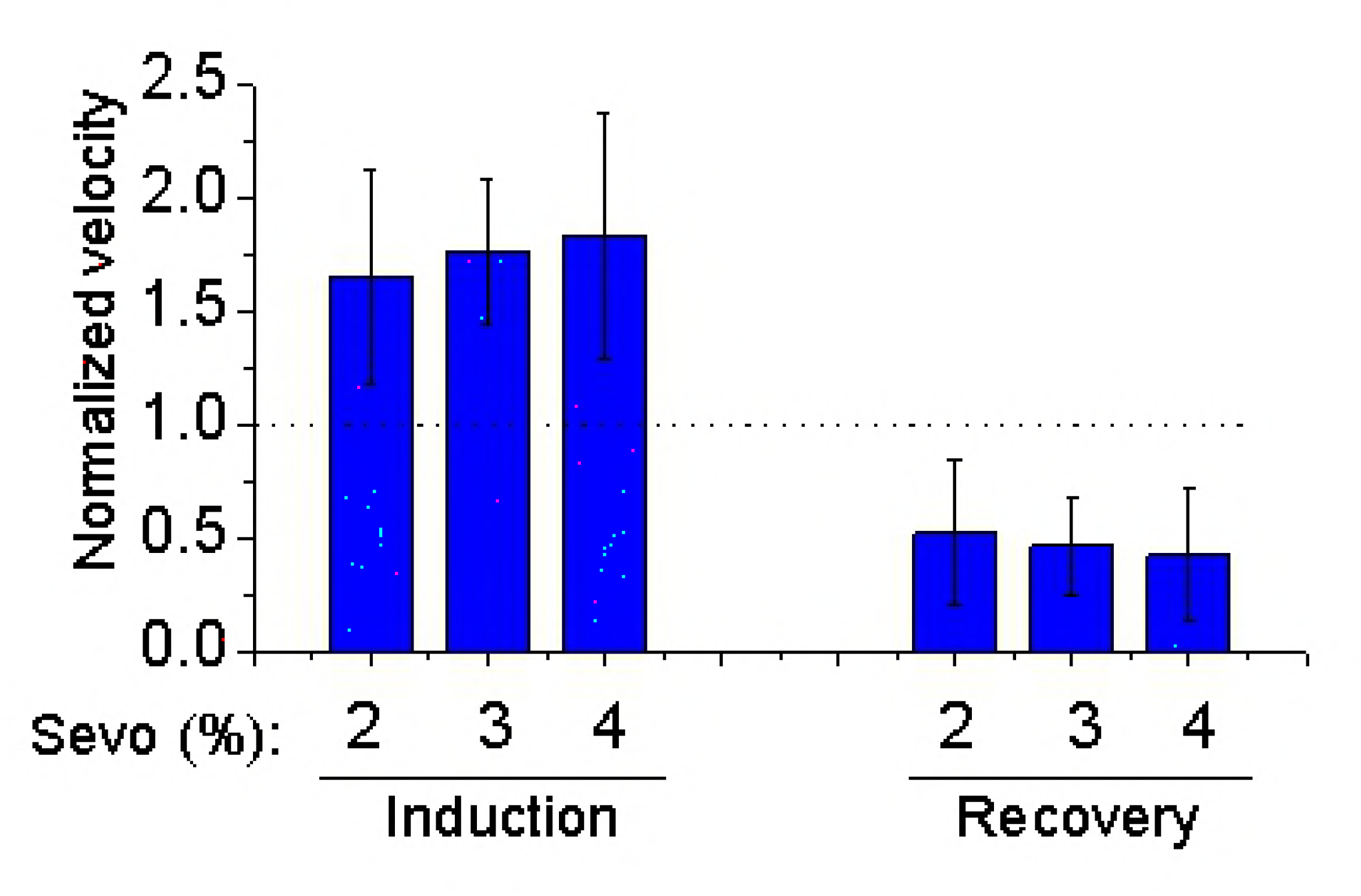
Ratio of mean speed induced by sevoflurane at different concentrations of sevoflurane and baseline. Normalized velocity shown as mean ±SEM using 4-6 mice in each group. Note that the sevoflurane-induced moving pattern (hyperactivity during induction and hypoactivity during recovery) in the mouse model was observed at all tested concentrations ranging from 2-4 % sevoflurane with undetectable statistical significance among the different concentrations.

### The sevoflurane-induced hyperactivity was greater in AD Tg mice

Recent *in vivo* evidence from mouse models and human patients have indicated that, particularly in early stages of AD, neuronal circuits are hyperactive instead of hypoactive^8–11^, which could lead to key complications in AD^12^. Therefore, we questioned how sevoflurane-induced movement in AD Tg mice.

To answer this question, we performed further experiments using AD Tg mice. Figure 3A shows that the sevoflurane-induced hyperactivity in AD Tg mice was greater than that of the WT mice (342.60 ± 42.43 cm/min for AD Tg mice versus 247.94 ± 36.83 cm/min for WT mice). Given the findings that the baseline velocity of mice was different, we calculated the ratio of locomotion of 2% sevoflurane to baseline levels in each mouse, graphing out the normalized parameters to show the results of locomotion in AD Tg mice and WT mice (Figures 3B and 3C). We found that the ratios of velocity and walking distance were still significantly higher in AD Tg mice as compared to those in WT mice (velocity: 3.30 ± 0.37 versus 1.76 ± 0.34; walking distance = 2.92 ± 0.47 versus 1.63 ± 0.48, p < 0.05, student’ s t-test). Taken together, these data showed that the exposure to sevoflurane increased the locomotion in WT and AD Tg mice, with a greater increase in the AD Tg mice.

**Figure 3.**
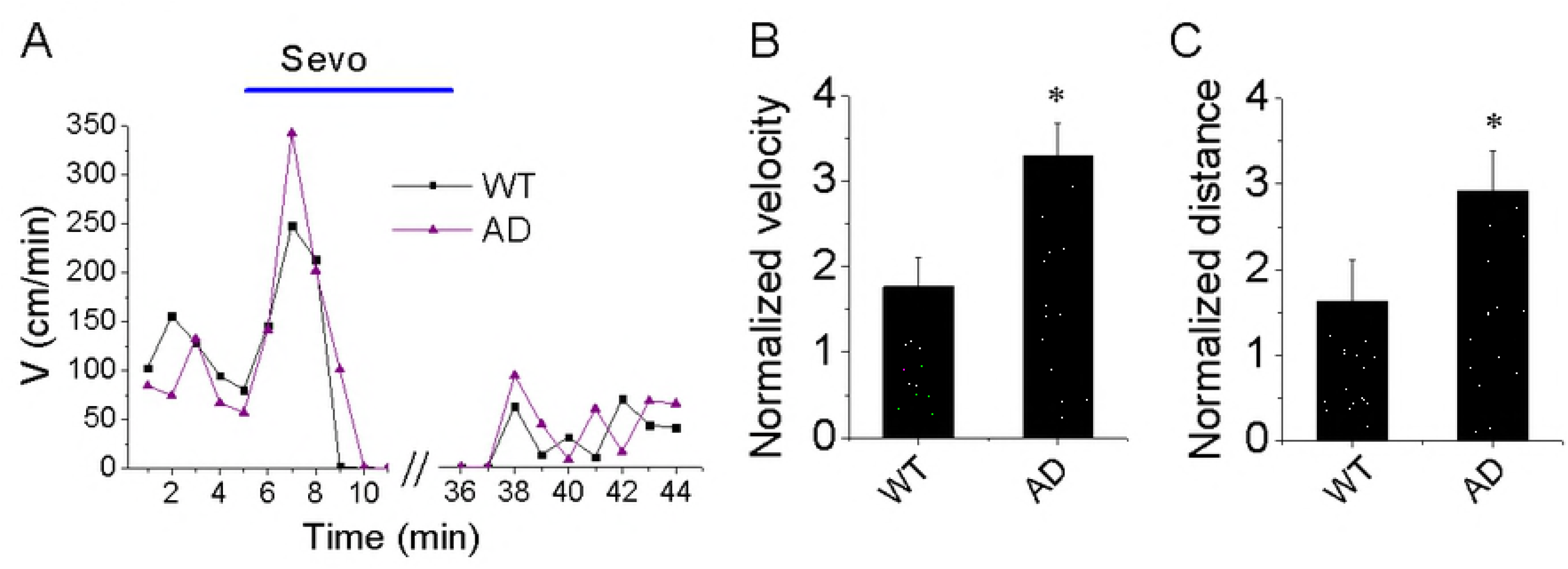
Sevoflurane-induced hyperactivity in AD Tg mice compared with WT mice. (A) The time-course of movement velocity in the activity chamber before and after the exposure to sevoflurane in the WT and AD Tg mice; (B&C) Summary of normalized velocity and walking distance in the induction period for each mouse type. Data shown as mean and SEM, n = 7-9 mice in each group. Note that the sevoflurane-induced hyperactivity is increased in the AD Tg mice.

### Sevoflurane did not change food intake and body weight of the mice

In a separate group of animals, we examined the effects of anesthesia with 3% sevoflurane for 4 hours on food intake and daily body weight in each of the mice. Figure 4 shows that the food intake and body weight of the mice in anesthesia-treated group were not statistically significant as compared to those observed in the mice in the control group from days 1 to 4 post anesthesia. These data demonstrate that the anesthetic sevoflurane did not significantly change the food intake and body weight of the mice.

**Figure 4.**
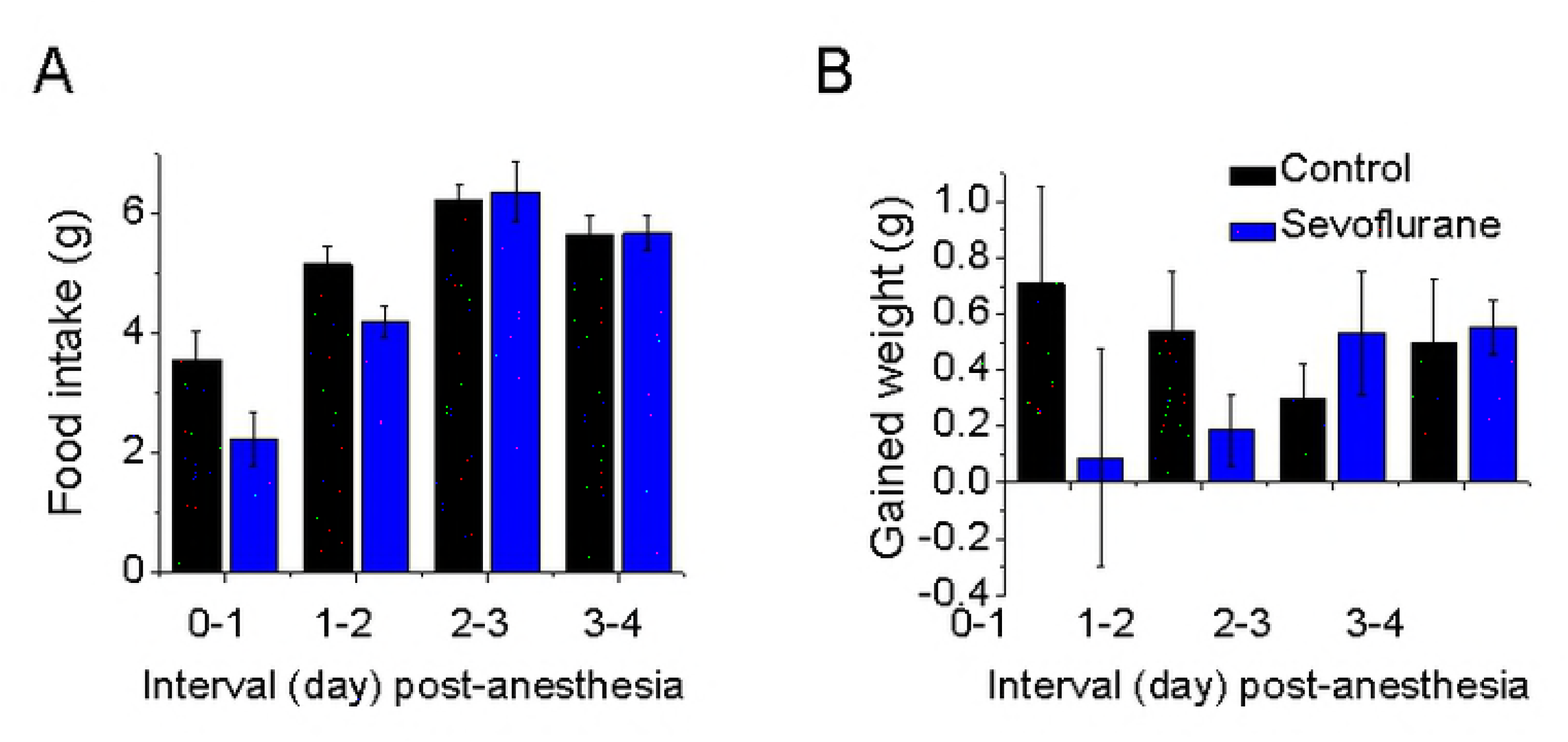
Food intake and weight gains after the 4-hour treatments of 3% sevoflurane. Data are showed as mean ± SEM from 14-15 mice in each group. Note that the statistically significant difference in food intake (A) and gained weight (B) from 1 to 4 days after the administration of sevoflurane was not observed between the mice in the sevoflurane-treated group and the mice in control group.

## Discussion

By measuring the velocity and walking distance at baseline levels, we examined the locomotion of the mice under sevoflurane-induced anesthesia. The data showed that sevoflurane, at clinically-relevant concentrations, induced hyperactivity during the induction phase and hypoactivity during the recovery phase of the anesthesia in the WT mice. Moreover, AD Tg mice showed greater hyperactivity following the sevoflurane anesthesia.

Our finding is also in line with the clinical observation that several general anesthetics, especially sevoflurane, induce emergence agitation and hyperactivity in patients ^10,12^. Moreover, previous studies showed sevoflurane at 1% had the ability to activate hippocampal CA3 kainate receptors to induce neuronal hyperactivity, which was measured by electrophysiology, during induction and recovery of the anesthesia ^16^. Whether the sevoflurane-induced activation of hippocampal CA3 kainate receptor contributes to the sevoflurane-induced hyperactivity, however, is not known. The future studies to test this hypothesis are warranted.

Moreover, the data showed that the sevoflurane-induced hyperactivity in AD Tg mice was greater than that observed in WT mice. Hyperactivity has been reported in patients with AD, which is associated with functional and structural alterations in a distributed network of brain regions supporting memory and other cognitive domains^17^. Evidence that brain hyperactivity occurs in early or pre-symptomatic AD has also been reported. Bookheimer et al., for example, noted that middle aged and elderly people who were carriers of the APOE epsilon4 allele, the genetic risk factor of AD, showed abnormally high activation in the hippocampus and other memory-related brain regions that are typically affected in AD during memory tasks^18^. Putcha et al. found hyperactivity in the hippocampus among patients with mild cognitive impairment^19–22^. Thus, the paradox is that higher levels of hippocampal activity may drive AD disease progress and may be early indicators of AD-related neurodegeneration^11,23^. Our data showed that sevoflurane induced greater hyperactivity in AD Tg mice than in WT mice. However, the exact the mechanism and the clinical relevance of such findings need to be further investigated.

Our study has several limitations. First, we did not demonstrate the concentration-dependent effects in the sevoflurane-induced hyperactivities. The exact reason is not clear at the present time. Our anesthetic delivery system did not allow the gas to immediately reach the expected concentrations (i.e. 2, 3 or 4%). Therefore, there was a time of delay (∼40s) which could prevent the observation of the possible concentration-dependent effect of sevoflurane-induced hyperactivity. Second, the data were obtained in mice, and the clinical relevance of the sevoflurane-induced hyperactivity in the mice remains to be determined.

In conclusion, we have provided the initial evidence that inhalation of the anesthetic sevoflurane induced an acute hyperactivity in the mice, particularly in the AD Tg mice. This novel observation of sevoflurane-induced hyperactivity would be helpful in the design of an animal model for future studies looking at emergence agitation and postoperative behavioral changes. Ultimately, these studies may promote further research into the mechanisms of anesthesia action and anesthesia neurotoxicity.

Author contributions
Hoai T. Ton, PhD.: This author designed, conducted the study, analyzed the data and wrote the manuscript.
Lei Yang, MD. MS.: This author conducted the study and analyzed the data.
Zhongcong Xie, MD, PhD.: This author designed the study and wrote the manuscript

## References

1. Vakkuri A, Yli-Hankala A, Särkelä M, et al. Sevoflurane mask induction of anaesthesia is associated with epileptiform EEG in children. ActaAnaesthesiolScand. 2001;45(7): 805–811. doi:10.1034/j.1399-6576.2001.045007805.x.

2. Liang P, Li F, Liu J, Liao D, Huang H, Zhou C. Sevoflurane activates hippocampal CA3 kainate receptors (Gluk2) to induce hyperactivity during induction and recovery in a mouse model. Br JAnaesth. 2017;119(5): 1047–1054. doi:10.1093/bja/aex043.

3. Dahmani S, Stany I, Brasher C, et al. Pharmacological prevention of sevoflurane- and desflurane-related emergence agitation in children: a meta-analysis of published studies. Br J Anaesth. 2010;104: 216–223. doi:10.1093/bja/aep376.

4. Yli-Hankala A, Vakkuri A, Särkelä M, Lindgren L, Korttila K, Jäntti V. Epileptiform electroencephalogram during mask induction of anesthesia with sevoflurane. Anesthesiology. 1999;91(6): 1596–1603. doi:10.1097/00000542-199912000-00009.

5. Pilge S, Jordan D, Kochs EF, Schneider G. Sevoflurane–induced Epileptiform Electroencephalographic Activity and Generalized Tonic-Clonic Seizures in a Volunteer Study. Anesthesiology. 2013;119(2): 447. doi:10.1097/ALN.0b013e31827335b9.

6. Julliac B, Cotillon P, Guehl D, Richez B, Sztark F. Target-controlled induction with 2.5% sevoflurane does not avoid the risk of electroencephalographic abnormalities. Ann Fr Anesth Reanim. 2013;32(10): e143–e148. doi:10.1016/j.annfar.2013.07.812.

7. Mohanram A, Kumar V, Iqbal Z, Markan S, Pagel PS. Repetitive generalized seizure-like activity during emergence from sevoflurane anesthesia. Can J Anesth Can d’ anesthésie. 2007;54(8): 657. doi:10.1007/BF03022961.

8. Stargardt A, Swaab DF, Bossers K. The storm before the quiet: neuronal hyperactivity and Aβ in the presymptomatic stages of Alzheimer’ s disease. NeurobiolAging. 2015;36(1): 1–11. doi:10.1016/J.NEUROBIOLAGING.2014.08.014.

9. Vossel KA, Tartaglia MC, Nygaard HB, Zeman AZ, Miller BL. Epileptic activity in Alzheimer’ s disease: causes and clinical relevance. Lancet Neurol. 2017;16(4): 311–322. doi:10.1016/S1474-4422(17)30044-3.

10. Bakker A, Krauss GL, Albert MS, et al. Neuron Report Reduction of Hippocampal Hyperactivity Improves Cognition in Amnestic Mild Cognitive Impairment. Neuron. 2012;74:467–474. doi:10.1016/j.neuron.2012.03.023.

11. Born HA. Seizures in Alzheimer’ s disease. Neuroscience. 2015;286:251–263. doi:10.1016/J.NEUROSCIENCE.2014.11.051.

12. Busche MA, Konnerth A. Neuronal hyperactivity - A key defect in Alzheimer’ s disease? BioEssays. 2015;37(6): 624–632. doi:10.1002/bies.201500004.

13. Lu Y, Wu X, Dong Y, Xu Z, Zhang Y, Xie Z. Anesthetic sevoflurane causes neurotoxicity differently in neonatal naïve and alzheimer disease transgenic mice. Anesthesiology. 2010;112(6): 1404–1416. doi:10.1097/ALN.0b013e3181d94de1.

14. Dong Y, Zhang G, Zhang B, et al. The common inhalational anesthetic sevoflurane induces apoptosis and increases β-amyloid protein levels. Arch Neurol. 2009;66(5): 620–631. doi:10.1001/archneurol.2009.48.

15. Ton HT, Phan TX, Abramyan AM, Shi L, Ahern GP. Identification of a putative binding site critical for general anesthetic activation of TRPA1. PNAS. 2017;114(14): 3762–3767. doi:10.1073/pnas.1618144114.

16. Vutskits L, Xie Z. Lasting impact of general anaesthesia on the brain: mechanisms and relevance. Nat Rev Neurosci. 2016;17(11): 705–717. doi:10.1038/nrn.2016.128.

17. Bookheimer SY, Strojwas MH, Cohen MS, et al. Patterns of brain activation in people at risk for Alzheimer’ s disease. NEngl JMed. 2000;343(7): 450–456. doi:10.1056/NEJM200008173430701.

18. Putcha D, Brickhouse M, O’ Keefe K, et al. Hippocampal Hyperactivation Associated with Cortical Thinning in Alzheimer’ s Disease Signature Regions in Non-Demented Elderly Adults. J Neurosci. 2011;31(48): 17680–17688. doi:10.1523/JNEUR0SCI.4740-11.2011.

19. Esmaeili MH, Bahari B, Salari A-A. ATP-sensitive potassium-channel inhibitor glibenclamide attenuates HPA axis hyperactivity, depression- and anxiety-related symptoms in a rat model of Alzheimer’ s disease. Brain Res Bull. 2018;137:265–276. doi:10.1016/j.brainresbull.2018.01.001.

20. Amatniek JC, Hauser WA, DelCastillo-Castaneda C, et al. Incidence and Predictors of Seizures in Patients with Alzheimer’ s Disease. Epilepsia. 2006;47(5): 867–872. doi:10.1111/j.1528-1167.2006.00554.x.

21. Hesdorffer DC, Hauser WA, Annegers JF, Kokmen E, Rocca WA. Dementia and adult-onset unprovoked seizures. Neurology. 1996;46(3): 727–730. doi:10.1212/WNL.46.3.727.

22. de Haan W, van Straaten ECW, Gouw AA, Stam CJ. Altering neuronal excitability to preserve network connectivity in a computational model of Alzheimer’ s disease. PLOS Comput Biol. 2017;13(9):e1005707. doi:10.1371/journal.pcbi.1005707.

23. Hauser WA, Morris ML, Heston LL, Anderson VE. Seizures and myoclonus in patients with Alzheimer’ s disease. Neurology. 1986;36(9): 1226–1230. doi:10.1212/WNL.36.9.1226.

